# RNA-xLSTM: Evaluating xLSTM as an Alternative Foundation to Transformers in RNA Modeling

**DOI:** 10.1101/2025.07.14.664653

**Authors:** Matija Pintarić, Rafael Josip Penić, Mile Šikić

**Affiliations:** Faculty of Electrical Engineering, University Of Zagreb, 3 Unska Street, Zagreb, 10000, Croatia; Genome Institute of Singapore (GIS), Agency for Science, Technology and Research (A*STAR), 60 Biopolis Street, Genome, Singapore, 138672, Republic of Singapore

## Abstract

Transformer-based architectures currently achieve state-of-the-art performance across a wide range of domains, including biological sequence modeling. Motivated by the recent introduction of the xLSTM architecture, we investigate its effectiveness for RNA sequence modeling by comparing a 33.7M-parameter RNA-xLSTM model against two leading RNA language models: RNA-FM and RiNALMo-33M. We pretrain RNA-xLSTM on the RNAcentral database and evaluate its performance on two downstream tasks: RNA secondary structure prediction and splice site prediction. Our results show that while RNA-xLSTM underperforms compared to the similarly sized RiNALMo, it does outperform the larger RNA-FM model on certain tasks. However, its overall performance remains inconsistent, and its advantages over transformer-based models are unclear, suggesting that further work is needed to assess its true potential in RNA modeling.

## 1 Introduction

The advent of the attention mechanism and the transformer architecture [1] significantly impacted the field of artificial intelligence, particularly in natural language processing. However, their influence extended into other domains, such as computational biology and genomics. One of the most famous examples of this is ESMFold [2], which leveraged attention-based mechanisms to achieve remarkable accuracy in protein structure prediction. This success resulted in an increased development of protein large language models.

Proteins are not the only beneficiaries of this paradigm shift. Recently, RNA molecules have also received attention from the scientific community. However, data for RNA is scarcer than for proteins, and is mostly unlabeled. To leverage this unlabeled data, several transformer-based RNA large language models have been introduced, including Uni-RNA [3], RNA-FM [4] and RiNALMo [5].

Although transformer models continue to dominate the landscape with state-of-the-art performance, alternative architectures, such as Mamba [6] and RWKV [7], have emerged to address their limitations in efficiency and scalability. One of the newest proposed architectures is the xLSTM [8], a modern evolution of the LSTM [9]. xLSTM introduces a novel exponential gating mechanism, an improved memory structure, and a reconfigured design that enables greater parallelizability.

Since its introduction, xLSTM has demonstrated strong performance across various domains, including computer vision [10] and biological sequence modeling, namely, Bio-xLSTM [11]. However, Bio-xLSTM solely focuses on DNA, protein, and chemical sequences, while its performance on tasks involving RNA sequences has not yet been explored. Motivated by this, we investigate the effectiveness of xLSTM on RNA-related tasks by comparing it with state-of-the-art RNA language models.

Following the approach in RiNALMo, we pretrain a 33.7M-parameter RNA-xLSTM model on the RNAcentral [12] database and evaluate it on downstream RNA-related tasks. We then compare its performance with two existing models: the 33.5M-parameter RiNALMo and the larger 99M-parameter RNA-FM. Specifically, we focus on two downstream tasks: RNA secondary structure prediction and splice site prediction. Our goal is to assess how RNA-xLSTM performs relative to these transformer-based models in terms of accuracy, generalization, and suitability for RNA sequence modeling.

## 2 Related work

### xLSTM

The main limitations of the standard LSTM include its inability to revise storage decisions, limited storage capacity, and lack of parallelizability. xLSTM addresses these limitations with two new cell types, the mLSTM and the sLSTM. The sLSTM cell introduces exponential gating for its input and forget gates. It also adds a normalizer and stabilizer state to prevent numerical overflow. The mLSTM cell builds on these changes by increasing the memory cell from a vector to a matrix and incorporating key and value projections, allowing memory retrieval via a covariance update rule [13]. It also abandons memory mixing to enable parallelization. The sLSTM and mLSTM cells are further integrated into specialized residual blocks. The sLSTM cell is placed into a residual block with post up-projection, while the mLSTM cell is integrated into a residual block with pre up-projection. The complete xLSTM model is formed by stacking residual sLSTM and mLSTM blocks, either separately or in combination.

### Bio-xLSTM

builds on the original xLSTM by introducing three domain specific variants: DNA-xLSTM, Prot-xLSTM and Chem-xLSTM, each available in multiple configurations, varying in block type and number, embedding dimension, context length and up-projection ratio. The key distinction among them lies in their pretraining tasks. We adapt the DNA-xLSTM variant as the basis for our model, using the same architectural design but with a slightly altered configuration, having more blocks and a larger embedding dimension. We focus exclusively on masked language modeling as the pretraining objective, in contrast to the original DNA-xLSTM, which uses both masked and causal language modeling. This choice is motivated by the fact that RNA shares a similar token vocabulary with DNA, making this variant the most natural fit for RNA sequences.

### RNA-FM

is a BERT-style transformer-based model pretrained on sequences from the RNAcentral database. It comprises of 12 transformer encoder blocks, each featuring a feed-forward layer with a hidden size of 640 and a multi-head self-attention mechanism with 20 heads, resulting in approximately 99 million parameters.

### RiNALMo

is a BERT-style transformer encoder. It is augmented with techniques such as Rotary Positional Embedding [14], the SwiGLU activation function [15] and FlashAttention-2 [16]. It is pretrained on non-coding RNA sequences from the RNAcentral, Rfam [17], nt [18], and Ensembl [19] databases. RiNALMo comes in several size varying configurations. We use the RiNALMo-33M variant, as it performs comparably or better than RNA-FM on several RNA related tasks, despite being nearly three times smaller. With 33.5 million parameters, it remains efficient to train and serves as a suitable baseline for models of this size. RiNALMo-33M consists of 12 transformer encoder layers, an embedding dimension of 480, 20 attention heads, and incorporates Rotary Positional Embeddings.

## 3 Methods

### 3.1 RNA-xLSTM

We use a configuration inspired by the DNA-xLSTM setup proposed in Bio-xLSTM, scaled to match the parameter count of RiNALMo-33M for a fair comparison. Specifically, we used the DNA-xLSTM-4M configuration, which uses only mLSTM blocks, and scale it up by increasing both the number of blocks and the embedding dimension. The final architecture consists of 15 mLSTM blocks, an embedding dimension of 600, Rotary Positional Embeddings, and a 2:1 up-projection ratio, resulting in a total of 33.7 million parameters, virtually matching the size of RiNALMo-33M.

We opted for a pure mLSTM configuration, as sLSTM blocks are not parallelizable during training and thus offer no practical advantage in our setting. Bidirectionality is achieved by flipping the input sequence after each block, following one of the approaches described in the Bio-xLSTM paper.

### 3.2 Data preparation

For the pretraining data preparation, we followed a pipeline nearly identical to that presented in RiNALMo to ensure the fairest possible comparison, with the only difference being that we used only the RNAcentral dataset. First, we downloaded the publicly available RNAcentral dataset. We then filtered out sequences shorter than 16 nucleotides and longer than 8192 nucleotides. Duplicate sequences were removed using *seqkit rmdup* [20], and the remaining sequences were clustered using *mmseqs easy-linclust* [21] with the command-line arguments *–min-seq-id 0*.*6* and *-c 0*.*8*. Clustering ensures that, during pretraining, each sequence in a batch is sampled from a different cluster at random, thereby maximizing diversity among the training examples. The resulting dataset was stored in LMDB (Lightning Memory-Mapped Database) format for faster access.

Additionally, we adopted the tokenization approach proposed in RiNALMo, using four primary tokens to represent the standard nucleotide bases: A, C, T, and G. For ambiguous or degenerate nucleotide codes, we included additional tokens: I, R, Y, K, M, S, W, B, D, H, V, N, and the gap symbol -. Special tokens were also incorporated into the vocabulary: a [CLS] token at the beginning of each sequence, a [EOS] token at the end, a [PAD] token for padding shorter sequences to a uniform maximum length, and a [MASK] token used during the pretraining objective.

### 3.3 Pretraining

Pretraining was conducted following the same approach as in RiNALMo. First, we use BERT-like

[22] masking of the training sequence. A total of 15% of the input tokens are selected for masking. Of these tokens, 80% are replaced with a [MASK] token, 10% are replaced with a random token, and 10% are left unchanged. The model was trained using cross-entropy loss computed only on the masked tokens.

Due to computational constraints, mainly GPU memory, input sequences were truncated to a maximum length of 1024 tokens (1022 nucleotides with [CLS] and [EOS] token). To maintain uniform length within each batch, any sequence shorter than 1,024 tokens was padded with the [PAD] token.

Pretraining was conducted on three A100 GPUs, each with 40 GB of memory, using a batch size of 140 and a total of 42,000 global steps. We used the AdamW [23] optimizer with weight decay set to 0.1. A linear warm-up scheduler was applied over the first 1,000 iterations, increasing the learning rate to 5 × 10^*™*4^. Following the warm-up phase, a cosine annealing schedule was used, with a minimum learning rate of 5 × 10^*™*5^. Gradient norms were clipped to a maximum value of 1.0 to ensure training stability.

To evaluate the quality of the embeddings obtained during pretraining and to compare the performance of RNA-xLSTM, we fine-tune and assess the model on two downstream tasks: RNA secondary structure prediction and splice site prediction. These tasks were also used in RiNALMo, both on RiNALMo-33M and RNA-FM models, providing a consistent and well established benchmark for comparison.

### 3.4 Secondary structure prediction

Secondary structure refers to the hydrogen bonds that form between bases when an RNA molecule folds in space. This structure plays a key role in how RNA functions, affecting its stability, interactions, and role in processes like gene regulation. Because of this, accurately modeling RNA secondary structure is a fundamental step in understanding RNA biology. The prediction of this structure is framed as a binary classification task, where for each pair of nucleotides in a sequence, the model predicts whether the nucleotides are paired or unpaired. We perform this task by attaching a prediction head to the pretrained RNA-xLSTM model. Specifically, we use a simple two-block ResNet [24] architecture as the prediction head. The output from the head is a matrix in which each element represents the probability logit that nucleotide pair *(i, j)* is paired. Since the pairing of *(i, j)* is equivalent to *(j, i)*, only the upper triangular part of the matrix above the main diagonal is included in the loss computation. This is the same head used in the RiNALMo paper for both RiNALMo and RNA-FM models, enabling a fair comparison. The training loss is binary cross-entropy.

We finetune and evaluate the models on two datasets: the bpRNA dataset from SPOT-RNA [25] and ArchiveII [26]. The bpRNA dataset comes pre-split into a training set (TR0), which also has a validation split, and a test set (TS0). However, it is important to note that sequences from the same RNA family can appear in both sets, making this dataset less suitable for evaluating generalization across families. To address this limitation, we also evaluate on the ArchiveII dataset, which contains 3,865 RNA sequences divided into nine distinct families. We use the approach where we train and validate on eight families, and evaluate on the unseen ninth. We repeat this for every family.

Following the approach in RiNALMo, we employ a layer unfreezing schedule during finetuning. For both datasets, only the prediction head is finetuned during the first three epochs. Subsequently, for bpRNA, the model is finetuned for an additional twelve epochs, and for ArchiveII, an additional three epochs. The number of epochs was selected based on optimal performance on the validation sets. For both datasets, the learning rate was initialized at 5 ×10^−4^ and linearly decreased to 5 × 10^−5^ over 7,500 steps.

Model performance is compared based on the average F1 score on the respective test sets. Additionally, following the approach in [27], predictions that are close enough are viewed as correct. A pair *(i, j)* is considered paired if either *(I* ± 1, *j)* or *(i, j* ± 1) is predicted as paired. The results from RiNALMo and RNA-FM were copied from the RiNALMo paper, where both of these models were finetuned and evaluated with this exact approach.

### 3.5 Splice site prediction

Splice site prediction is a sequence-level classification task where the model determines whether a RNA sequence contains a donor or acceptor splice site. These sites indicate the boundaries between exons and introns. Accurate prediction of splice sites is important for understanding how RNA is processed before translation and can provide insights into gene regulation and potential splicing-related disorders.

We evaluate two approaches for this task. In the first, the representation of the [CLS] token is passed to the prediction head. In the second, we compute the average pooling over all token embeddings in the sequence and use this as input to the prediction head. The motivation for exploring the average pooling approach stems from architectural differences: unlike RNA-FM and RiNALMo, which inherently support bidirectional context through self-attention, RNA-xLSTM achieves bidirectionality by alternating the input direction across layers. As a result, relying solely on the [CLS] token may limit its ability to capture global sequence context effectively. The prediction head is the same one as proposed in RiNALMo, to achieve fair comparison between the models. It consists of a two layer multilayer perceptron with a hidden size of 128, with a GELU activation function. Cross-entropy is used as the training loss.

The model is finetuned on the GS_1 dataset from [28]. We finetune it separately for both the donor and acceptor tasks. Finetuning was performed for three epochs, with a batch size of 32. Learning rate was fixed at 1 ×10^−5^. During finetuning, we perform a 10 fold cross-validation to find the optimal number of epochs for the finetuning. The testing is done on four different unseen organisms: Danio, Fly, Thaliana and Worm. The models are compared based on the averaged F1-score on these organisms. Same as for the secondary structure task, the results from RNA-FM and RiNALMo are copied from the RiNALMo paper, where both models were finetuned and evaluated with the approach described here.

## 4 Results

### 4.1 Pretraining

To monitor the training progress, we tracked the perplexity metric on the validation set over the course of 42,000 global steps. As shown in Figure 1, the validation perplexity consistently decreased from the maximum possible value of 22, indicating stable convergence of the model during pretraining.

**Figure 1.**
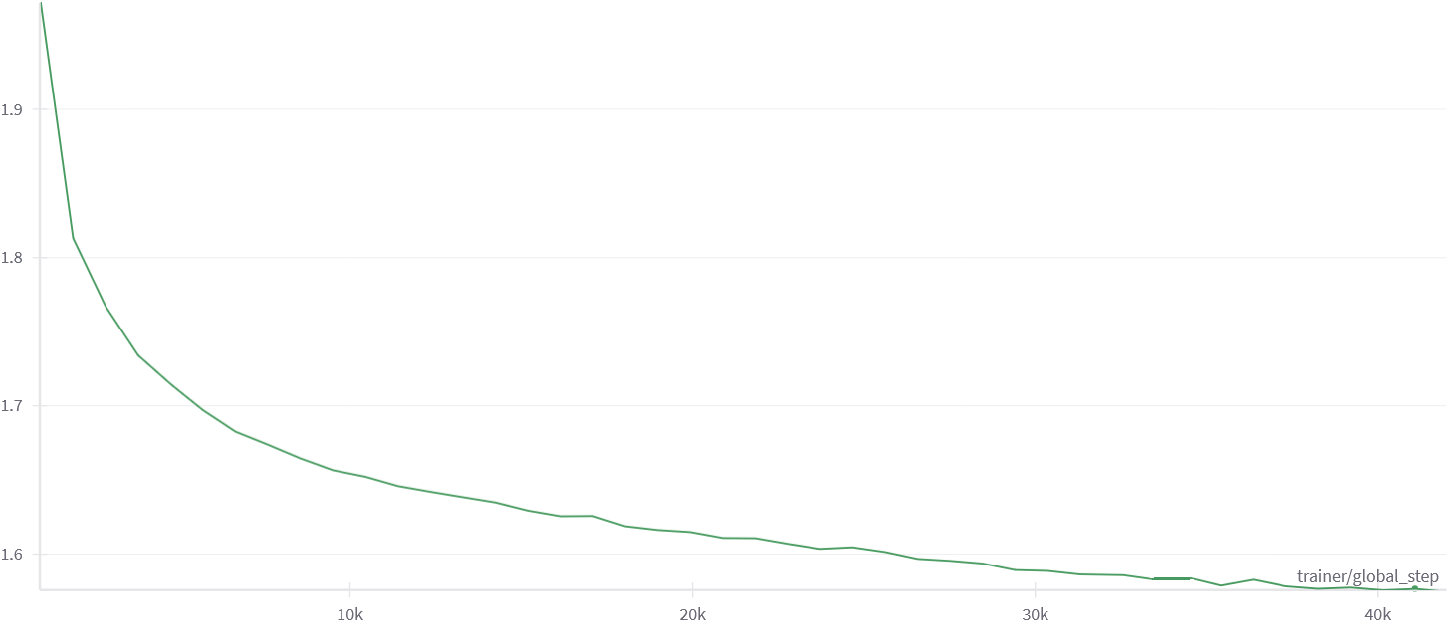
Validation perplexity curve over 42,000 steps during pretraining.

### 4.2 Secondary structure prediction

We first performed fine-tuning on the bpRNA dataset. The results are presented in Table 1. Based on the F1 scores, the RNA-xLSTM model performs comparably to RNA-FM, even slightly better, while falling short of RiNALMo. However, this comparison should be interpreted with caution, as the bpRNA training and test sets are not split by RNA families. As a result, the model’s ability to generalize to unseen families cannot be reliably assessed.

**Table 1:**
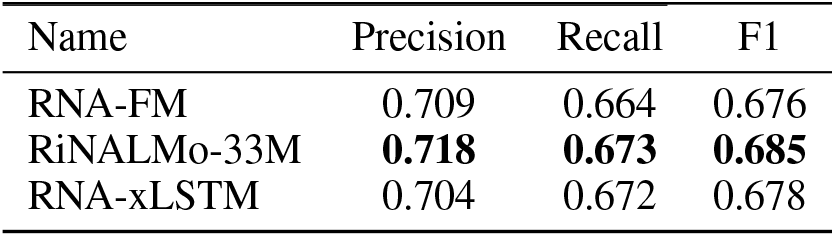
F1 score of secondary structure prediction on the bpRNA dataset

A more robust evaluation of the generalization capabilities is provided by the ArchiveII dataset, which was evaluated on a family unseen in training. The results of this approach for each family are presented in Table 2.

**Table 2:**
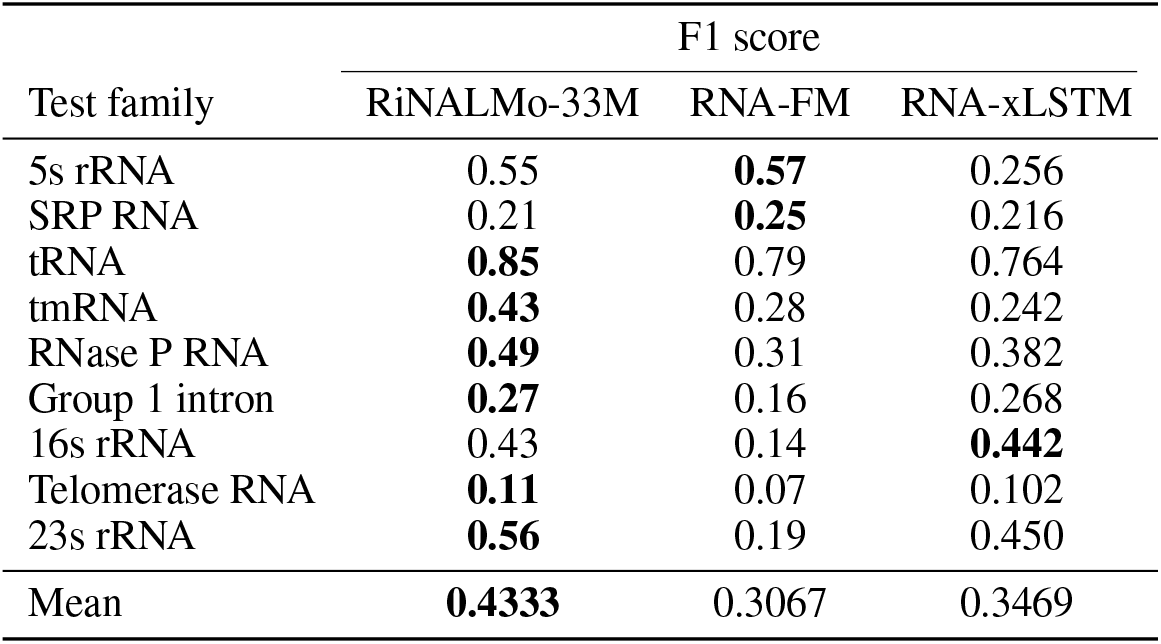
F1 score for each test family in the ArchiveII dataset for the secondary structure prediction task

| Test family | F1 score |  |  |
| --- | --- | --- | --- |
|  | RiNALMo-33M | RNA-FM | RNA-xLSTM |
| 5s rRNA | 0.55 | **0.57** | 0.256 |
| SRP RNA | 0.21 | **0.25** | 0.216 |
| tRNA | **0.85** | 0.79 | 0.764 |
| tmRNA | **0.43** | 0.28 | 0.242 |
| RNase P RNA | **0.49** | 0.31 | 0.382 |
| Group 1 intron | **0.27** | 0.16 | 0.268 |
| 16s rRNA | 0.43 | 0.14 | **0.442** |
| Telomerase RNA | **0.11** | 0.07 | 0.102 |
| 23s rRNA | **0.56** | 0.19 | 0.450 |
| Mean | **0.4333** | 0.3067 | 0.3469 |

Overall, RiNALMo achieves the highest mean F1 score. Interestingly, RNA-xLSTM ranks second, outperforming the significantly larger RNA-FM model, which has nearly three times as many parameters. Notably, RNA-xLSTM, like RiNALMo and RNA-FM, struggles with the telomerase RNAs, achieving a low F1 score on this subset. This family contains the longest sequences in the dataset, raising the question of whether current language models can effectively capture long-range dependencies required for accurate secondary structure prediction. This is particularly noteworthy in the case of the xLSTM architecture, which has been positioned as a strong alternative to transformers for modeling long range interactions. This observation highlights the importance of further investigating how different architectures, including xLSTM, handle long-range dependencies in RNA, and suggests a valuable direction for future work.

### 4.3 Splice site prediction

The results of splice site prediction task are presented in Table 3.

**Table 3:**
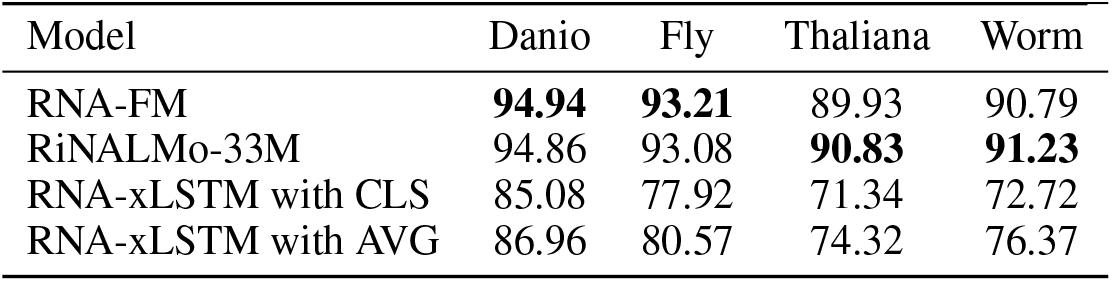
F1 score on splice site prediction task on four different test sets

| Model | Danio | Fly | Thaliana | Worm |
| --- | --- | --- | --- | --- |
| RNA-FM | **94.94** | **93.21** | 89.93 | 90.79 |
| RiNALMo-33M | 94.86 | 93.08 | **90.83** | **91.23** |
| RNA-xLSTM with CLS | 85.08 | 77.92 | 71.34 | 72.72 |
| RNA-xLSTM with AVG | 86.96 | 80.57 | 74.32 | 76.37 |

RNA-xLSTM underperformed relative to both RNA-FM and RiNALMo across all test sets, regardless of whether the [CLS] token or average pooling was used. However, the results suggest that average pooling offers a more effective representation for splice site prediction, likely due to its ability to capture broader contextual information than the [CLS] token alone. This raises an important question about whether the xLSTM architecture can capture global context as effectively as transformer-based models. While xLSTM is designed to handle long-range dependencies, these findings indicate that its current implementation may not fully leverage this strength in sequence level classification tasks.

## 5 Conclusion

We pretrained a 33.7M parameter RNA-xLSTM model, evaluated it on two downstream tasks, and compared its performance with state-of-the-art transformer-based models for RNA sequence modeling. While RNA-xLSTM demonstrates promising results on the RNA secondary structure prediction task, occasionally outperforming larger transformer models, it still does not consistently match the best performing approaches. Its performance on the splice site prediction task is notably weaker, which may point to challenges in capturing global context using its current architecture, particularly when relying on the [CLS] token for sequence level predictions.

Although adjustments such as improved tokenization strategies may help close the performance gap, it remains uncertain whether xLSTM can fully match transformer-based models on RNA sequence modeling tasks. These results suggest that while xLSTM has potential, further testing on a wider range of downstream tasks is needed to assess whether xLSTM can meaningfully challenge the performance of current transformer architectures.

## 6 Code availability

The code repository is available at code repository and the pre-trained weights are available at model weights. We also provide scripts for finetuning the pre-trained model on the downstream tasks. The data for the downstream tasks can automatically be downloaded using the scripts provided in the repository.

## Acknowledgements

The authors would like to thank Ivona Martinović and Tin Vlašić for their insights and comments for this work.

